# Temporal profiling of CD4 T-cell activation and differentiation upon SARS-CoV-2 spike protein immunisation

**DOI:** 10.1101/2022.07.21.500987

**Authors:** José Almeida-Santos, Rita Berkachy, Chanidapa Adele Tye, Jehanne Hassan, Bahire Kalfaoglu, Murray E. Selkirk, Masahiro Ono

## Abstract

CD4 T-cells require T-cell receptor (TCR) signalling for their activation and differentiation. Foxp3+ regulatory T-cells (Treg) are dependent on TCR signals for their differentiation and suppressive function. However, it is not fully known how TCR signalling controls the differentiation of polyclonal CD4 T-cells upon antigen recognition at the single-cell level in vivo. In this study, using Nr4a3-Tocky (<u>T</u>imer-<u>o</u>f-<u>c</u>ell-<u>k</u>inetics-and-activit<u>y</u>), which analyses temporal changes of antigen-reactive T-cells following TCR signalling, we investigated T-cell response to Spike protein fragments (S1a, S1b, S2a, and S2b) upon immunisation. We show that S1a and S2a induced the differentiation of PD1^hi^CXCR5^+^ T follicular helper (Tfh) cells, which is related to CD4 T-cell immunogenicity. In contrast, S1b induced CD25^hi^GITR^hi^PD-1^int^ Treg, which intermittently received TCR signalling. Using Foxp3-Tocky, which analyses *Foxp3* transcriptional dynamics, the S1b-reactive Treg sustained *Foxp3* transcription over time, which is a hallmark of activated Treg. Foxp3 fate-mapping showed that the S1b-reactive Treg were derived not from pre-existing thymic Treg, suggesting Foxp3 induction in non-Treg cells. Thus, the current study reveals temporally dynamic differentiation of CD4 T-cells and Treg upon immunisation in the polyclonal TCR repertoire.

## INTRODUCTION

T-cells play central roles for protection against intracellular pathogens and in vaccination, including severe acute respiratory syndrome coronavirus 2 (SARS-CoV-2), which causes Coronavirus Disease 2019 (COVID-19) (*1*). Typically, primary T-cell responses are delayed primarily because antigen-specific T-cells are rare - estimated to be 10 to 1000 cells per epitope, due to the enormous diversity of T-cell receptor (TCR) repertoire (i.e. polyclonal T-cells) in physiological conditions (*2-4*). However, upon infection or vaccination, antigen-specific T-cells receive TCR signals, which triggers their activation and proliferation, leading to their differentiation into mature T-cell subsets and expansion.

While CD8 T-cells eliminate infected cells by recognising viral antigens, CD4 T-cells coordinate immune responses through helper and regulatory functions. Antigen recognition in CD4 T-cells promotes the differentiation of T follicular helper (Tfh) cells in the context of vaccination. Tfh cells express high levels of the checkpoint molecules PD-1 and ICOS together with the chemokine receptor CXCR5 (*5*), which is used by Tfh for its recruitment to the germinal centre (GC) in lymphoid follicles to help B-cell maturation (*6*). In addition, CD4 T-cells help CD8 T-cells (i.e. T-cell priming) to induce T-cell memory (*7*). In addition, some antigens may induce regulatory T-cells (Treg), which suppress B-cell and T-cell responses, especially in the intestines (*8*). In contrast, the suppressive counterpart of Tfh, T follicular regulatory (Tfr) cells, can prevent T-cell help to B-cells (*9, 10*). Tfr cells can differentiate from pre-existing Treg cells (*11*), most of which are reactive to either self-antigen or microbiome antigen in healthy mice and humans (*8, 12*). However, this may be dependent on antigen or disease model, as Tfh cells can upregulate Foxp3 during the end-stage of GC reactions (*13*).

It remains unclear how TCR signalling dynamically controls the activation and differentiation of polyclonal CD4 T-cells to antigen in vivo. Current methods for analysing antigen-specific T-cells in the polyclonal T-cell repertoire are limited and include MHC-tetramer, transgenic TCR mouse lines, and retroviral gene transduction of TCR into primary T-cells (*14, 15*). Nur77-GFP transgenic reporter mice have been a major tool to investigate TCR signal strength and antigen-reactive T-cells (*16*). However, due to the long half-life of GFP (∼ 56 hours (*17*)), the use of an MHC-tetramer is still required for the analysis of antigen-specific T-cells (*18*). However, MHC-tetramer staining of CD4 T-cells is complicated due to non-specific binding of Class II MHC-tetramers to LAG-3 on the surface of activated CD4 T-cells (*19*). It therefore remains an outstanding question as to whether and how immunological properties of antigens, or T-cell immunogenicity, can control CD4 T-cell activation and differentiation. T-cell immunogenicity is typically addressed by cytokine and activation marker analysis upon in vitro stimulation of T-cells using each epitope (*20*). However, these assays cannot provide insights into how individual antigen-specific T-cells are activated and differentiate in vivo.

Successful vaccines against COVID-19 use the spike (S) protein of SARS-CoV-2, which induces a protective immunity through neutralising antibodies and memory T-cells (*21, 22*). Whilst neutralising antibodies target the receptor-binding domain (RBD) and N-terminal domain (NTD) of S protein (*23, 24*), unsurprisingly, T-cell epitopes for CD4 are found throughout the protein in humans with some enrichment in NTD and RBD in humans (*25, 26*) and in mice (*27*). Importantly, SARS-CoV-2 can infect mice especially when the virus is adapted to mice (*28, 29*). In addition, S-protein immunisation can induce protective T-cell memory in mice (*30*). These support the relevance of using mice to analyse T-cell responses against SARS-CoV-2. However, it is not clear what types of CD4 T-cell subsets are induced by S-protein immunisation and whether different domains of S protein can induce different CD4 T-cell subsets.

In this study, we analysed T-cell response to the S-protein of SARS-CoV-2 using <u>T</u>imer-<u>O</u>f-<u>C</u>ell-<u>K</u>inetics-and-activit<u>Y</u> (Tocky). Using the fluorescent Timer protein (Timer), which spontaneously and irreversibly shifts its emission spectrum from blue fluorescence (Blue) to red fluorescence (Red), Tocky allows analysis of temporally dynamic changes of individual cells in vivo. Nr4a3-Tocky is a Timer reporter mouse line for the TCR signal downstream gene *Nr4a3*, can be used to analyse antigen-reactive T-cell responses and differentiation (*31-33*) by showing temporal changes in T-cell activation and differentiation following TCR signals in vivo (*34*). Thus, in this study, we aimed to analyse S protein-induced polyclonal T-cell response using Nr4a3-Tocky and to understand antigen-dependent control of T-cell differentiation and the properties of S protein in relation to T-cell immunogenicity in vivo.

## RESULTS

### Nr4a3-Tocky allows analysis of antigen-reactive T-cells by their TCR reactivity

We produced the following 4 fragments of the S-protein of SARS-CoV-2: S1a (14-312) containing the N-terminal domain; S1b (313-680) containing the RBD; S2a (681-911) containing the fusion peptide and S2b (912-1212) containing the two heptapeptide repeat sequences, HR1 and HR2 (**Fig. 1A**). To analyse the properties of each fragment for inducing CD4 T-cell activation and differentiation, we investigated temporally dynamic T-cell response by immunising Nr4a3-Tocky mice with these four S protein fragments. Mice were challenged with each antigen one week later and draining lymph node (DLN) cells were analysed at 24h following the challenge (**Fig. 1B**).

**Figure 1.**
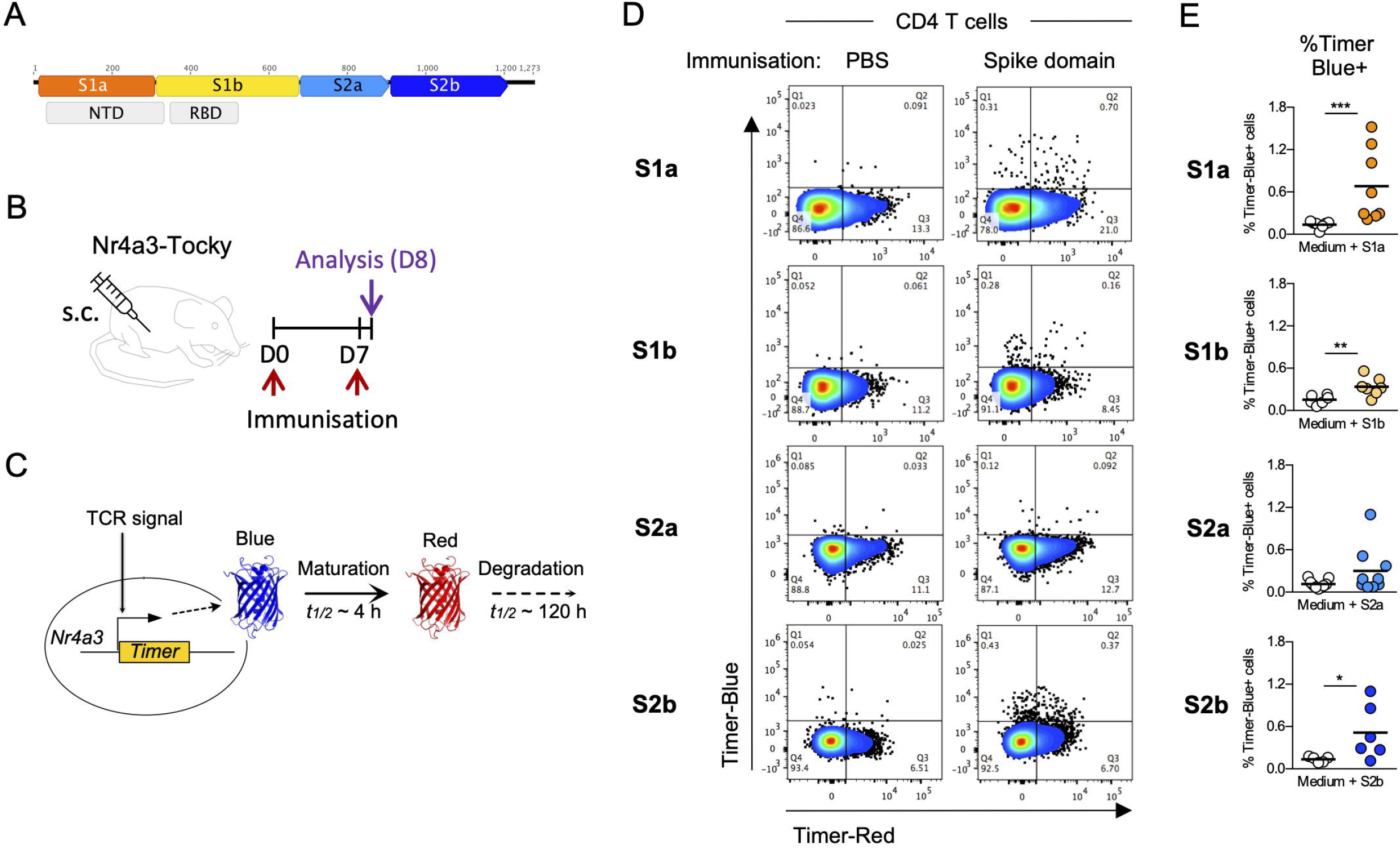
Nr4a3-Tocky allows analysis of antigen-reactive T-cells by their TCR reactivity. **(A)** Scheme depicting each of the SARS-CoV-2 spike protein fragments “domains” overlaid with the N Terminal (NTD) and the receptor binding domain (RBD). **(B**) Scheme depicting the immunisation regimen used for each fragment. Mice were immunised with either 1.5 μg of S fragment or PBS emulsified in Incomplete Freund’s adjuvant (IFA) containing 50 μg of poly(I:C) and challenged 7 days later in absence of added adjuvant. Analysis took place 24h later. (**C)** Diagram showing the induction and maturation of the Timer protein over time upon TCR signalling. TCR signals activate Nr4a3 Timer transcription and its translation results into immature Timer protein with blue fluorescence (Blue). The immature chromophore then spontaneously and irreversible matures, changing its fluorescence to red (Red). Maturation half-life of the Timer protein is 4 hours while mature protein is stable with a half-life > 120 hours. (**D)** Draining lymph node cells (DLN) from mice immunised as in B) were incubated 48h in the presence of the corresponding immunisation fragment without exogenous TCR stimulation. Shown is the percentage of Nr4a3-Timer Blue^+^ CD4 T cells in each sample after the incubation period. (**E**) Scatter plots summarising data in (D). For each protein fragment tested, data are pooled from 2 independent experiments (n = 2-5) except for S2a, pool of 3 independent experiments (n =2-4). Statistical analysis performed using nonparametric Mann-Whitney test. *P < 0.05, **P < 0.005, and ***P < 0.001.

First, we tested if Nr4a3-Tocky allows identification of antigen-specific T-cells to each of the S-protein fragments. DLN cells from Nr4a3-Tocky mice with or without immunisation with one of the four fragments were incubated *in vitro* for 48h with the corresponding S-protein fragment and analysed by flow cytometry. Since Timer expression is induced specifically by TCR signalling in Nr4a3-Tocky (*34*) and its Blue fluorescence decays with the half-life of 4 hours (*35*), Blue+ indicates that T-cells have recognised their cognate antigen and received TCR signalling in the culture (**Fig. 1C**). Whilst CD4 T-cells from PBS immunised mice incubated with each S-protein fragment scarcely expressed Blue, Blue was induced in CD4 T-cells from mice immunised with S1a, S1b, and S2b, but not S2a (**Fig. 1D, 1E**). This indicates that S-fragment specific CD4 T-cells were expanded by the immunisation and can respond to the rechallenge in vitro. S1a, S1b, and S2b are therefore likely to be more T-cell immunogenic than S2a in mice.

### S1a immunisation induces antigen-reactive Tfh cells with unique TCR signal dynamics

Next, we performed ex-vivo flow cytometric analysis to analyse effects of S1a immunisation on CD4 T-cells. Nr4a3-Tocky mice were immunised with the S1a fragment (or PBS for control, with poly(I:C) as adjuvant) at day 0 and challenged at day 7 (by S1a or PBS). CD4 T-cells from DLN were analysed by flow cytometry 24 hours after challenge (**Fig. 2A**). S1a immunisation did not change the percentage of Timer^+^ cells within CD4 T-cells or that of Foxp3^+^ CD4 T-cells (**Fig. S1A – S1C**), suggesting that S1a-reactive T-cells are rare. Next, we tested if S protein immunisation changed the composition of antigen-reactive T-cells as identified by Nr4a3-Tocky. However, as expected, PD-1^hi^ CXCR5^+^ Tconv cells were increased by S1a immunisation (**Fig. 2B**). Paradoxically, S1a immunisation decreased the percentage of Timer^+^ cells in PD-1^hi^ CXCR5^+^ Tconv cells **(Fig. 2C, 2D)**.

**Figure 2.**
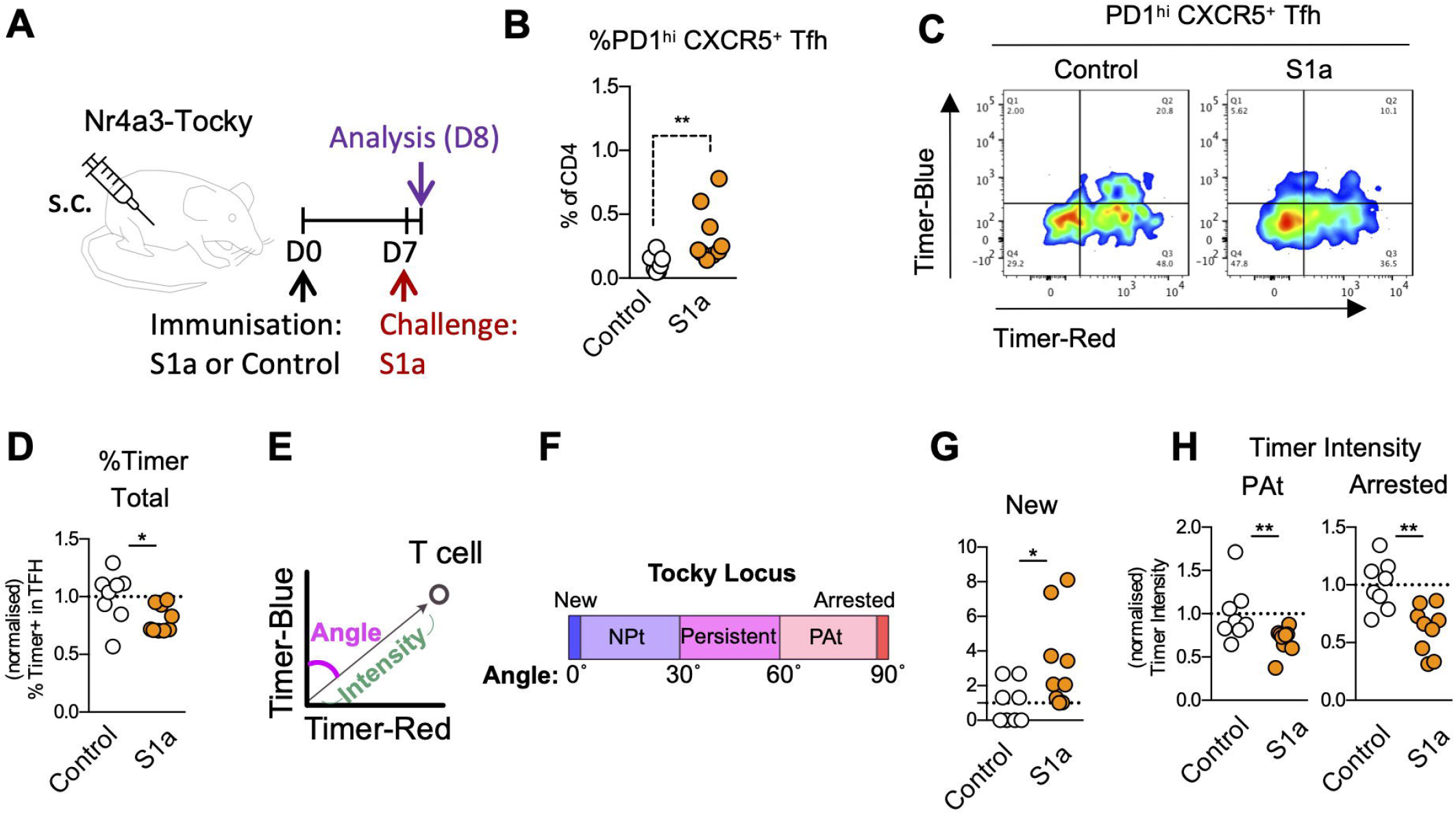
S1a immunisation induces antigen-reactive Tfh cells with unique TCR signal dynamics. **(A)** Scheme depicting the immunisation regimen. Mice were immunised with either 1.5μg of S1a fragment or PBS emulsified in Incomplete Freund’s adjuvant (IFA) containing 50μg of Poly(I:C) and challenged 7 days later in absence of added adjuvant. Analysis took place 24h later. **(B)** Percentage of all Tconv PD1^hi^ CXCR5^+^ cells in the total CD4 T cell population. **(C-D)** Nr4a3-Blue and -Red expression in Tconv PD1^hi^ CXCR5^+^ cells (C) and their normalised percentage of total Nr4a3-Timer^+^ cells (D). **(E-F)** Angle transformation and Tocky locus approach. Trigonometric transformation for Timer Angle and Intensity (E). Timer Angle is categorised as follows: New (Angle 0°), NPt (0° - 30°), Persistent (Pers, 30° - 60°), PAt (60° - 90°) and Arrested (Arr, 90°) (F). **(G-H)** Timer locus analysis of PD-1^hi^ CXCR5^+^ Tconv cells. Shown are the normalised percentage of cells in the New locus (G) and the normalised values for Timer Intensity in the Persistent to Arrested (PAt) and Arrested locus (H). Data from two independent experiments (n = 4-5). Statistical analysis performed using nonparametric Mann-Whitney test and parametric t-test *P < 0.05, **P < 0.005.

To investigate the T-cell dynamics underlying the S1a effect, Tocky analysis was applied to the flow cytometric data. Here Timer fluorescence data from Nr4a3-Tocky mice are quantitatively analysed by converting Blue and Red data into Timer Angle and Intensity, which is defined as angle from the Blue axis and distance from Timer(-) cells in the Blue-Red plane, respectively (**Fig. 2E** (*34*)). Angle data can be quantitatively analysed further by defining the Tocky locus: New (Angle 0°), New-to-Persistent transition (NPt, 0° - 30°), Persistent (30° - 60°), Persistent-to-Arrested transition (PAt; 60° - 90°) and Arrested (90°) (**Fig. 2F** (*34*)). T-cells that have recognised cognate antigen recently will be found in the New locus. T-cells will be found in the Persistent locus when they persistently engage with antigen. Antigen-experienced T-cells without active antigen engagement will be found in the Arrested locus.

Interestingly, S1a immunisation increased frequency of Tfh cells in the New locus (**Fig. 2G**), indicating that a higher rate of Tfh cells responded to the second immunisation than control. Given the decrease of Timer+ Tfh cells, S1a-reactive Tfh cells may have reduced Timer proteins most probably due to proliferation. In fact, S1a immunisation reduced Timer Intensity in those cells in the PAt and Arrested loci **(Fig. 2H, S1D**), indicating that less Timer proteins were accumulated in those Tfh cells.

Next, we tested if any additional novel population were induced by S1a immunisation. To this end, we applied a multidimensional Tocky analysis, which seamlessly analyses Angle and Intensity with marker expression data using the dimensional reduction methods Principal Component Analysis (PCA) and Uniform Manifold Approximation and Projection (UMAP) (*33*).

UMAP analysis showed that T-cells with low Angle were partially overlapped with those expressing high CD25 and CD69 expression, but not those with high PD-1 and CXCR5 expression (**Fig. S1E**). Computational clustering did not show any increased clusters, although identifying a significantly decreased cluster, Cluster 6 (**Fig. S1F, S1G**). UMAP analysis of Foxp3+ Treg did not show significant changes (**Fig. S1H – S1J**). This suggests that S1a-reactive T-cells did not differentiate into a unique T-cell populations, apart from the Tfh population, within a week after the initial immunisation.

### S1a booster immunisation induces antigen-reactive Tfh and transitional T-cells

Next, we tested if repeated S1a immunisation enhances the differentiation of S1a-reactive T-cells *in vivo*. Nr4a3-Tocky mice were immunised with S1a at days 1 and 15 (“boost”), while the control group was immunised with only one dose at day 1 (“single”). Both groups were challenged with the S1a fragment 2 weeks later after the last dose, and DLN were analysed at 24 hours after the challenge (**Fig. 3A**). As expected, PD-1^hi^ CXCR5^+^ Tfh cells were further increased by the booster dose of S1a (**Fig. 3B, 3C**). The additional boost dose did not increase the percentage of Timer+ within total CD4 T-cells, whether in Tconv or Treg (**Fig. S2A**).

**Figure 3.**
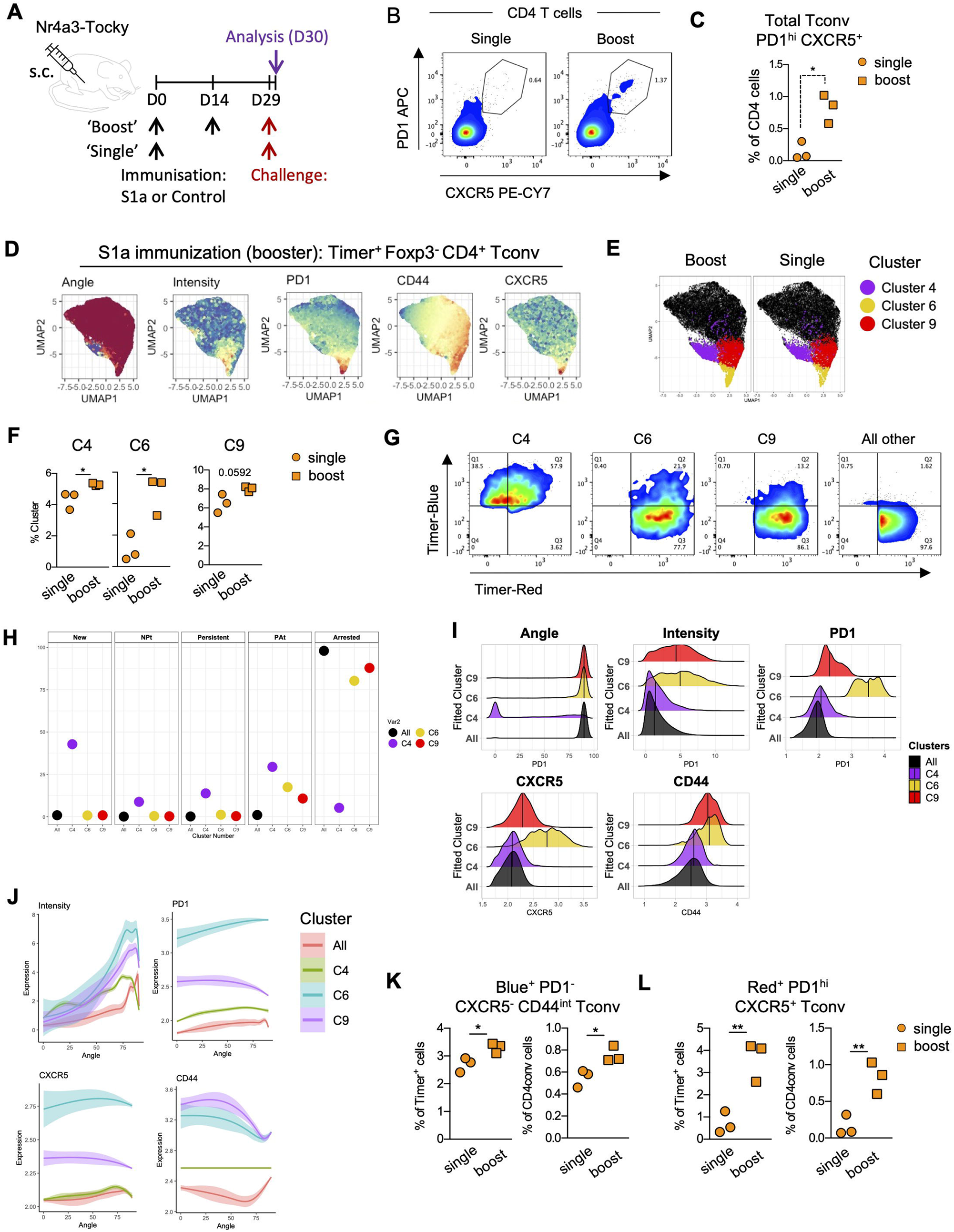
S1a immunisation induces antigen-reactive Tfh cells and transitional population upon booster immunisation. **(A)** Scheme depicting the immunisation regimen. Mice were immunised with 1.5μg of S1a fragment and after two weeks received either boost S1a (Boost) or PBS (Single). Two weeks later both groups were challenged with S1a and analysis took place 24h later. **(B-C)** Shown are the PD-1 and CXCR5 expression in total CD4 T cells (B) and percentage of Tconv PD-1^hi^ CXCR5^+^ in total CD4 T cells (C). **(D)** Uniform Manifold Approximation and Projection (UMAP) analysis of Tconv Timer^+^ cells. Heatmaps show Timer Angle and Intensity as well as the expression of the indicated markers. The UMAP was applied to Tconv Timer^+^ cell data from draining lymph nodes (DLN) of both S1a and PBS immunised mice. UMAP was performed using Timer-Blue, Timer-Red, CD25, CD44, PD-1, and CXCR5 as input data. **(E)** Clusters 4, 6 and 9 identified by k-means clustering overlaid in the UMAP space from (D). (**F)** Shown are the percentages of Tconv Timer^+^ cells in Clusters 4,6 and 9 identified in (E) (n = 4). **(G-J)** Cluster 4,6 and 9 were characterised. Shown are their Nr4a3-Blue and -Red expression (G), the percentage of cells in each Timer Locus (H), their Angle, Intensity and expression of PD-1, CXCR5 and CD44 (I) and the dynamical expression of the indicated markers along the Timer angle (J). **(K)** Percentage of Timer Blue^+^ PD-1^-^ CXCR5^-^ CD44^int^ cells in Tconv Timer^+^ (Left) and total Tconv cells (Right) from mice receiving single vs boost S1a immunisation (n = 3). **(L)** Percentage of Timer Red^+^ PD-1^hi^ CXCR5^+^ cells in Tconv Timer^+^ (Left) and total Tconv cells (Right) from mice receiving single vs boost S1a immunisation (n = 3). Statistical analysis performed using nonparametric Mann-Whitney test and parametric t-test *P < 0.05, **P < 0.005.

UMAP analysis showed that Intensity was correlated with the expression of PD-1 and CXCR5 (**Fig. 3D**). Computational clustering identified 9 clusters of cells (**Fig. S2B**). Tconv cells in Cluster 4 and Clusters 6 were significantly increased by the S1a boost, whilst those in Cluster 9 showed a marginal increase (p = 0.059) (**Fig. 3E - 3F, S2C)**. Most T-cells in Cluster 4 were Blue^+^, suggesting that Cluster 4 Tconv cells were enriched with antigen-experienced S1a-specific T-cells, which promptly responded to the challenge (**Fig. 3G**). Strikingly, 43% of Cluster 4 cells were in the New locus (**Fig. 3H**), indicating that UMAP captured T-cells in the ultra-early phase of activation, within 4 hours after T-cell antigen recognition (*35*). Clusters 6 and 9 Tconv cells were predominantly in the Arrested locus (80% and 88%, respectively) and also in the PAt locus (17% and 10%, respectively). Most cells in the other clusters were in the Arrested locus (98%, ‘All others’, **Fig. 3H**). This indicates that Clusters 4, 6, and 9 encompass the majority of antigen-reactive Tconv cells that were actively engaged with antigen.

Cluster 6 expressed the highest levels of Timer intensity PD-1 and CXCR5, which was followed by Cluster 9 expressing the second highest levels of these markers (**Fig. 3H**), This indicates that Cluster 6 Tconv cells are the most mature Tfh cells in the dataset. CD44 expression was high in both Clusters 6 and 9. Given the proximity of Cluster 9 to Cluster 6 in UMAP (**Fig. 3E**) and the Timer Angle progression (**Fig. 3H**), Cluster 9 Tconv cells are likely to be a transitional population differentiating into Tfh cells. Cluster 6 Tconv cells the most remarkably sustained Intensity and PD-1 expression as Angle increased (**Fig. 3I, Fig. S2C**), suggesting that the Tfh phenotype is dependent on TCR signals. Cluster 9 Tconv cells displayed increased Timer Intensity as Timer Angle increased, peaking at the PAt locus, which was similar to Cluster 6 Tconv cells. The expression of PD-1 and CXCR5 in Cluster 9 Tconv cells was lower than Cluster 6 and higher than Cluster 4 all others, across Angle (**Fig. 3J, Fig. S2C**). Lastly, we used a manual gating approach to identify Clusters 4 and 6-equivalent populations. We found that the booster dose increased the Cluster 4 equivalent Blue^+^ PD-1^lo^ CXCR5^-^ CD44^int^ Tconv and the Cluster 6 equivalent Red^+^ PD-1^hi^ CXCR5^+^ Tfh cells (**Fig. 2K, 2L**).

Collectively, these results suggest that the repeated S1a immunisation allowed antigen-reactive T-cells to differentiate into the Cluster 6 mature Tfh population, via the transitional populations Clusters 4 and 9.

### S1b immunisation induces Tfh and CD25^hi^GITR^hi^PD1^int^ Treg population

Next, we investigated the effects of S1b immunisation on CD4 T-cells. Using the same regimen for S1a (**Fig. 2A**), Nr4a3-Tocky mice were immunised with S1b fragment at day 0, challenged at day 7, and mice were analysed 24 hours after the challenge (**Fig. 4A**).

**Figure 4.**
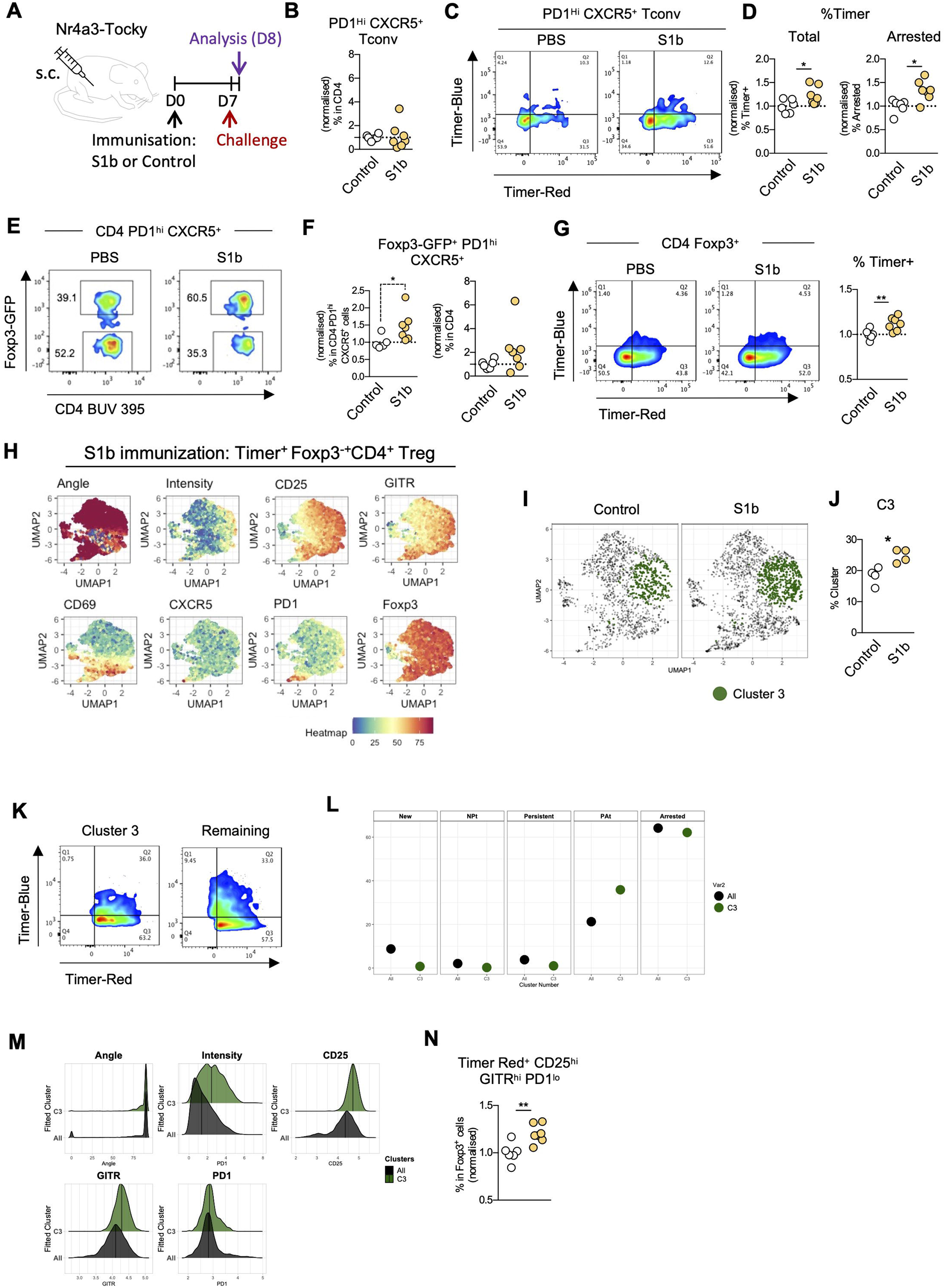
S1b immunisation induces Tfh and CD25^hi^GITR^hi^PD-1^int^ Treg population. **(A)** Mice were immunised with either S1b fragment or PBS emulsified in Incomplete Freund’s adjuvant (IFA) containing 50 μg of poly(I:C) and challenged 7 days later in absence of added adjuvant. Analysis took place 24h later. **(B)** Normalised percentage of Tconv PD-1^hi^ CXCR5^+^ cells within the total CD4 T cell population. **(C)** Nr4a3-Blue and -Red expression in Tconv PD-1^hi^ CXCR5^+^ cells. **(D)** Timer locus analysis of Tconv PD-1^hi^ CXCR5^+^ cells from mice immunised with either S1b or PBS, showing the normalised percentages of total Timer^+^ cells (Left) and cells in the Arrested locus (Right). **(E-F)** Analysis of Foxp3^+^ PD-1^hi^ CXCR5^+^ cells from mice immunised in (A). Shown are the expression of Foxp3-GFP in total PD-1^hi^ CXCR5^+^ cells (E) and the normalised percentage of Foxp3^+^ PD-1^hi^ CXCR5^+^ cells within the total PD-1^hi^ CXCR5^+^ fraction (Left) and within the total CD4 population (Right) (F). Data from 2 independent experiments (N=2-4) **(G)** Nr4a3-Blue and -Red expression and percentage of total Timer^+^ cells in CD4 Foxp3^+^ cells. **(H)** UMAP analysis of CD4 Foxp3^+^ Timer^+^ cell data from mice described in (A). UMAP was performed using Timer-Blue, Timer-Red, Foxp3, CD25, CD44, GITR, OX40, PD-1, CXCR5, and CD69 as input data. Heatmaps show Timer Angle and Intensity as well as the expression of the indicated markers. (**I)** Cluster 3 identified by k-means clustering overlaid in the UMAP space from (H). **(J)** Percentage of CD4 Foxp3^+^ Timer^+^ cells in Cluster 3 from either S1b or PBS immunisation samples (N=4). **(K)** Nr4a3-Blue and -Red expression of CD4 Foxp3^+^ Timer^+^ cells in Cluster 3 and in all concatenated clusters except Clusters 3 and 6. **(L)** Percentage of cells in each Timer Locus from Cluster 3 and all concatenated Clusters. **(M)** Histograms showing the Timer Angle and Intensity as well as the expression of the indicated markers in Cluster 3 and in all concatenated clusters. **(N)** Normalised percentage of Timer Red^+^ CD25^hi^ GITR^hi^ PD-1^int^ cells in total CD4 Foxp3^+^ from mice immunised with either S1b or PBS. Data from 2 independent experiments (N=2-4). Statistical analysis performed using nonparametric Mann-Whitney test. *P < 0.05, **P < 0.005.

First, we investigated the PD-1^hi^ CXCR5^+^ fraction of T-cells. Unlike S1a, S1b immunisation did not increase the frequency of Timer+ cells or PD-1^hi^ CXCR5^+^ Tfh cells in Tconv (**Fig. S3A, Fig. 4B**). UMAP analysis of antigen-reactive Tconv did not identify any significantly increased or decreased cluster (**Fig. S3B**-**D**). However, S1b immunisation increased the percentage of Timer+ cells in the PD-1^hi^ CXCR5^+^ Tfh, which was mostly due to the increase of Tfh cells in the Arrested locus (**Fig. 4C, 4D**), suggesting that the proportion of S1b-reactive cells increased within the PD-1^hi^ CXCR5^+^ Tfh fraction. Notably, the percentage of Foxp3+ cells in the PD-1^hi^ CXCR5^+^ fraction was increased, although the percentage of PD-1^hi^ CXCR5^+^ Foxp3^+^ T_FR_ cells was not significantly increased in CD4 T-cells (**Fig. 4E, 4F**). These results suggest that, although the expansion of S1b-specific T-cells was moderate, the composition of the PD-1^hi^ CXCR5^+^ fraction was changed and more enriched with S1b-specific T-cells, especially PD-1^hi^ CXCR5^+^ Foxp3^+^ T_FR_ cells. S1b did not change the percentage of Timer+ cells or the Timer profile of PD-1^hi^ CXCR5^+^ Foxp3^+^ T_FR_ (**Fig. S3E, S3F**).

Notably, the percentage of Timer+ cells in the total Foxp3^+^CD4 T-cells was significantly increased, although the effect was moderate (**Fig. 4G**). To identify S1b-reactive Treg population in the multidimensional space, we performed UMAP analysis of Nr4a3-Timer^+^ Treg cells (**Fig. 4H**). Computational clustering identified 7 clusters (**Fig. S3G**), of which Cluster 3 only was significantly increased by S1b immunisation (**Fig. 4I - 4J, Fig. S3H**). Cluster 3 cells had a higher percentage of cells in the PAt locus than the remaining cells (36% vs 21%) while Cluster 3 cells had few cells in the New, NPt, and Persistent loci (**Fig. 4K, 4L**), indicating their uniquely intermittent TCR signal dynamics. Cluster 3 Treg also highly expressed CD25 and GITR while showing an intermediate PD-1 expression (**Fig. 4M**). We used a manual gating approach and confirmed that Red^+^ CD25^hi^ GITR^hi^ PD-1^int^ Treg were significantly increased by S1b immunisation (**Fig. 4N**). Finally, we tested if the booster S1b immunisation would expand any of those Treg populations (**Fig. S3I**). However, unlike S1a, the booster dose did not enhance the expansion of any Tfh or Treg population, including PD-1^hi^ CXCR5^+^ Tfh and T_FR_ cells (**Fig. S3J**).

### S1b immunisation activates *Foxp3* transcription in S1b-reactive Treg

Next, we investigated Foxp3 transcriptional dynamics in response to S1b immunisation, using Foxp3-Tocky mice. As reported before, the bacterial artificial chromosome (BAC) transgenic *Foxp3*^*Timer*^ reports Foxp3 transcriptional activities by Timer protein in the Foxp3-Tocky model (*35*). Foxp3-Tocky mice were immunised with either S1b or PBS following a challenge (S1b or PBS) at 1-week after the immunisation (**Fig. 5A**). As expected, the majority of Timer+ cells in both groups showed a Blue^low^ Red^hi^ profile, which indicates that *Foxp3* transcriptional activities are intermittent in the majority of Treg (**Fig. 5B, 5C**). While the frequency of the total Timer^+^ cells in CD4 T-cells was similar between the two groups, S1b immunisation increased Treg cells in the Foxp3-Persistent locus but decreased those in the PAt and Arrested loci (**Fig. 5D**). In line with this result, the mean Timer Angle was significantly reduced by S1b immunisation despite no changes in Timer intensity (**Fig. 5E**). These collectively indicate that S1b immunisation activated *Foxp3* transcription in some Foxp3-Tocky^+^ Treg.

**Figure 5.**
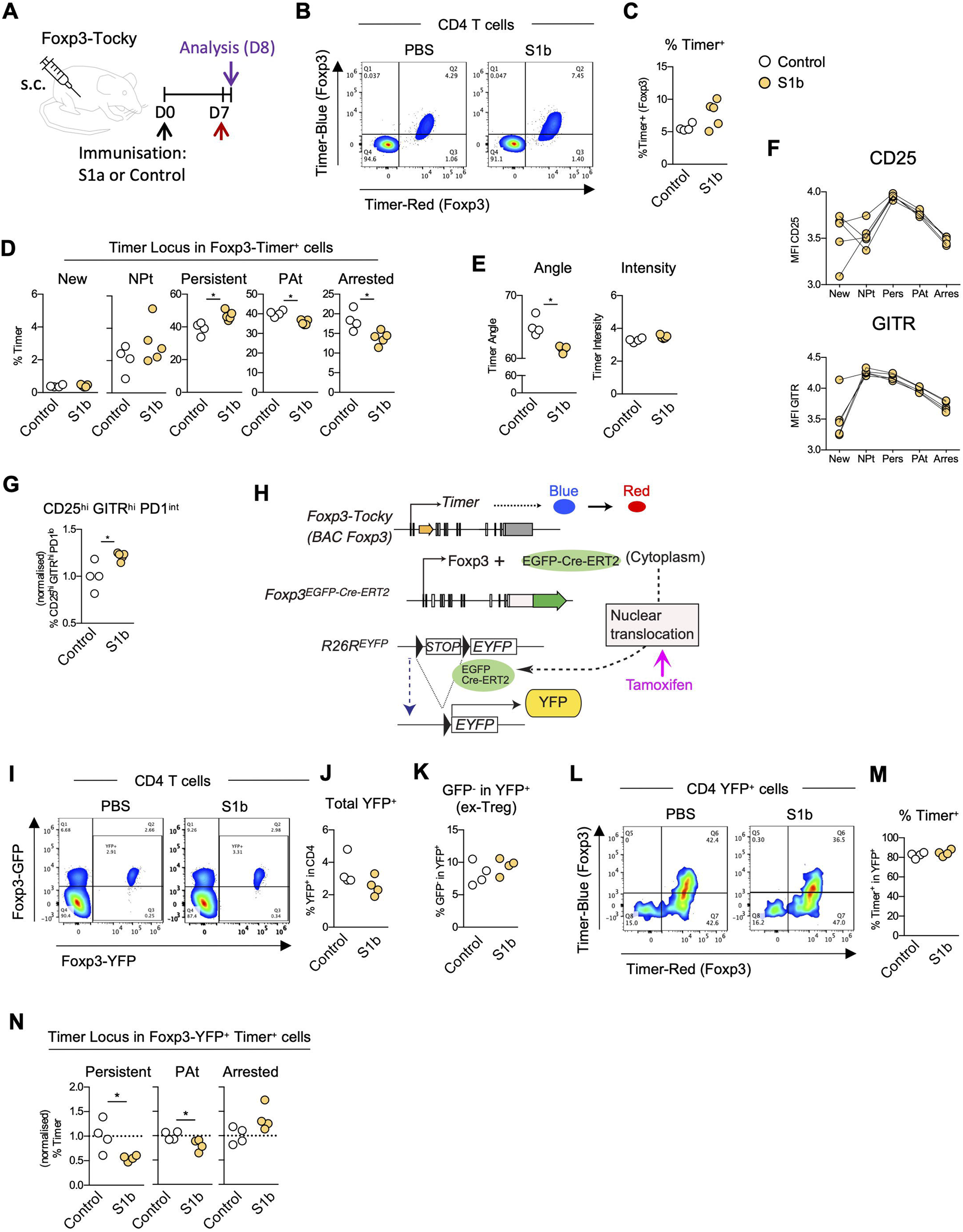
S1b immunisation activates Foxp3 transcription in S1b-reactive Treg. **(A)** Foxp3-Tocky Mice were immunised with either S1b fragment or PBS emulsified in Incomplete Freund’s adjuvant (IFA) containing 50μg of Poly(I:C) and challenged 7 days later in absence of added adjuvant. Analysis took place 24h later. Data from 2 independent experiments (N=1-3). **(B-E)** Shown are the expression of Foxp3 Timer-Blue and –Red (B) and the percentage of Timer^+^ cells in the CD4 population (C), the percentage of cells in the indicated Foxp3-Timer locus (D), and the Angle and Intensity of Foxp3-Timer (E). **(F)** Mean Fluorescence Intensity (MFI) of CD25 (top) and GITR (bottom) in each Foxp3-Timer locus in samples from mice immunised with S1b. (**G)** Normalised percentage of CD25^hi^ GITR^hi^ PD-1^int^ cells within total Foxp3-Timer^+^ cells of mice immunised as in (A). **(H-N)** Foxp3^CRE-YFP^ x Foxp3-Timer (Fate Mapper Foxp3-Timer) mice were administered tamoxifen (2 mg/mouse) intraperitoneally (i.p.) for 3 consecutive days. 4 days later, mice were immunised s.c. with either S1b or PBS and both 1 week and 2 weeks later received boost immunisation in absence of added adjuvant. 24h after the last immunisation DLN cells were analysed. Data from 2 independent experiments (n = 2). Shown are a schematic representation of the Fate Mapper Foxp3-Tocky system (H), the expression of Foxp3-GFP and Foxp3-YFP in CD4 T cells (I), the percentage of labelled YFP^+^ (J) and GFP^-^YFP^+^ (ex-Treg) (K) cells in the CD4 population, the expression of Foxp3 Timer-Blue and -Red (L) and the percentage of Timer^+^ cells in the labelled CD4 YFP^+^ fraction (M), and the percentage of cells in the indicated Foxp3-Timer locus of CD4 YFP^+^ cells (N). Statistical analysis performed using nonparametric Mann-Whitney test. *P < 0.05, **P < 0.005.

Using Nr4a3-Tocky mice our UMAP analysis had revealed that the S1b reactive Treg fraction was enriched in antigen-reactive CD25^hi^ GITR^hi^ PD-1^int^ T-cells The mean fluorescence intensity (MFI) of CD25 and GITR was the highest at the Persistent and NPt loci, respectively (**Fig. 5F**), indicating that sustained *Foxp3* transcription upregulates CD25 and GITR expression in the immunisation model, similar to a skin allergy model as reported previously (*35*). As expected, S1b immunisation increased the percentage of CD25^hi^ GITR^hi^ PD-1^int^ T-cells within Foxp3-Tocky^+^ CD4 T-cells (**Fig. 5G**), repeating the result by Nr4a3-Tocky (**Fig. 4N**). By integrating the results from Nr4a3-Tocky and Foxp3-Tocky, upon the challenge, S1b-reactive Treg receive TCR signals, as demonstrated by Nr4a3-Tocky, sustain *Foxp3* transcription as shown by Foxp3-Tocky, increasing CD25 and GITR expression, but not PD-1.

To address if S1b-reactive Treg were enriched with pre-existing Treg (thymic Treg), we developed the triple transgenic mouse line *Foxp3*^*EGFP-cre-ERT2*^ *R26R-EYFP Foxp3-Tocky* (designated as *Foxp3-Tocky::Foxp3 fate-mapping mice*). Here *Foxp3*^*EGFP-cre-ERT2*^ express the fusion protein EGFP-Cre-ERT2 under the endogenous Foxp3 promoter (*36*). The *R26R-EYFP* locus expresses YFP once Cre excises the floxed STOP cassette (*37*). Thus, Tamoxifen induces a constitutive YFP expression in Treg cells which are actively transcribing *Foxp3* at the time of the administration. The use of Foxp3-Tocky allows identification of antigen-reactive Treg as persistent *Foxp3* expressor as shown above (**Fig. 5H**).

Foxp3-Tocky::Foxp3 fate-mapping mice were administered with tamoxifen for 3 successive days (d1, d2, and d3) and immunised with either S1b (or PBS for control) on days 7 and 14. Then, mice were challenged with S1b (or PBS for control) on day 21 and analysed at 24 hours after the challenge. S1b immunisation did not change the percentage of YFP-expressing Treg (**Fig. 5I, 5J**). Since the efficiency of YFP labelling is around 30% of Treg, the Foxp3 fate mapping allows analysis of YFP+, but not YFP-in a meaningful manner (*36*). The percentage of YFP^+^GFP^-^, ‘ex-Treg’, was similar between the two groups (**Fig. 5K**). S1b immunisation did not change the percentage of the total Foxp3-Timer^+^ in YFP+ cells (**Fig. 5L, 5M**). However, Treg cells in the Persistent and PAt loci were reduced in S1b immunised group, compared to the control (**Fig. 5N**). Given that S1b-reactive Treg are found in the Persistent and PA-t loci of Foxp3-Tocky (**Fig. 5D**), this result suggests that S1b-reactive Treg are enriched with newly differentiated Treg upon immunisation.

### S2b domain induces antigen-reactive Tfh cells

Next, using Nr4a3-Tocky, we investigated antigenic properties of the remaining of S-protein fragments, S2a and S2b (**Fig. S4A**). S2a immunisation did not show any effects on the percentage of Tconv PD-1^hi^ CXCR5^+^ cells (**Fig. S4B**). S2a immunisation only marginally reduced the percentage of Timer^+^ cells within Tconv PD-1^hi^ CXCR5^+^ cells, and it did not change the percentage of Timer^+^ cells within either Tconv or Treg cells (**Fig. S4C-D**). UMAP analysis did not identify any S2a-reactive cluster in both Tconv and Treg (**Fig. S4E-H**). These results suggest that S2a is poorly immunogenic to mouse CD4 T-cells, whether Tconv or Treg.

S2b immunisation (**Fig. 6A**) did not change the overall accumulation of antigen-reactive Tconv cells (**Fig. S4I - S4J**). UMAP analysis of Tconv cells identified 8 clusters (**Fig. 6B, S4K**) and showed that S2b immunisation increased Cluster 1 (**Fig. 6C-D, S4L**). Cluster 1 cells were enriched with cells in the PAt locus (**Fig. 6E, 6F**) and highly expressed CD44, PD-1 and CXCR5 (**Fig. 6G**). In fact, S2b immunisation increased PD-1^hi^ CXCR5^+^ Tfh (**Fig. 6H**). Thus, S1a and S2b can induce Tfh cells. However, in contrast to S1a, S2b immunisation did not change the percentage of Nr4a3-Tocky^+^ cells within the Tfh fraction (**Fig. 6I-6J**), although it reduced Timer intensity in Tfh (**Fig. 6K**). S2b did not have any significant effects on Foxp3+ Treg by conventional and UMAP analysis (**Fig. S4M-S4N**).

**Figure 6.**
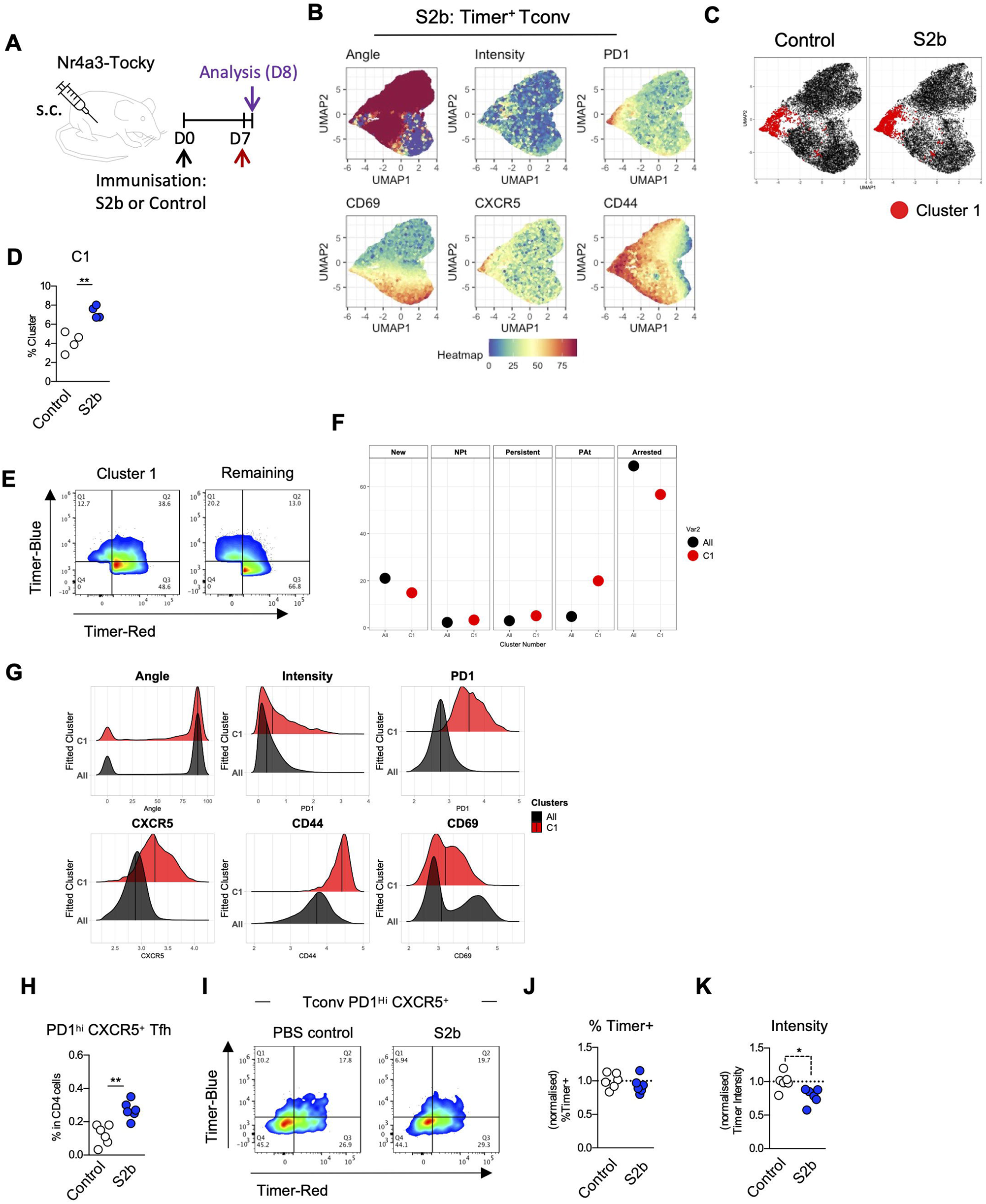
S2b domain induces antigen-reactive Tfh cells. **(A)** Mice were immunised with either S2b fragment or PBS emulsified in Incomplete Freund’s adjuvant (IFA) containing 50μg of poly(I:C) and challenged 7 days later in absence of added adjuvant. Analysis took place 24h later. **(B)** Uniform Manifold Approximation and Projection (UMAP) analysis of Tconv Timer^+^ cells from Nr4a3-Tocky mice immunised with either S2b or PBS. The UMAP was performed using Timer-Blue, Timer-Red, CD25, CD44, GITR, PD1, CXCR5, and CD69 as input data and the heatmaps show Timer Angle and Intensity as well as the expression of the indicated markers. **(C)** Cluster 1 identified by k-means clustering overlaid in the UMAP space from (B). **(D)** Percentage of Tconv Timer^+^ cells in Cluster 1 from either S2b or PBS samples (n=4). **(E)** Nr4a3-Blue and -Red expression of Tconv Timer^+^ cells in Cluster 1 and in all remaining concatenated clusters. **(F)** Percentage of cells in each Timer Locus from Cluster 1 and all concatenated Clusters. **(G)** Overlaid histograms showing Timer intensity and the expression of the indicated markers in Cluster 1 and all concatenated Clusters identified in (C). **H-K)** Analysis of total Tconv PD1^hi^ CXCR5^+^ cells from mice immunised as in (A). Shown are the percentage of the analysed cells within the total CD4 T cell population (H), their Nr4a3-Blue and -Red expression (I) and percentage of total Timer^+^ cells (J) and Timer Intensity in total Timer^+^ cells (K). Statistical analysis performed using nonparametric Mann-Whitney test. *P < 0.05, **P < 0.005.

## DISCUSSION

Currently, S-protein reactive T-cells have been identified and defined by the induction of activation markers (e.g., OX-40, 4-1BB and CD69) and/or the production of cytokines (e.g., interleukin-2, interferon-γ) upon peptide-stimulation in humans and mice (*27, 38*). However, it is unclear if these assays can identify non-cytokine producing antigen-reactive T-cell populations, especially Treg, because Treg show activated phenotype and do not produce most cytokines (*39, 40*). Here Nr4a3-Tocky provides a unique solution to identify antigen-reactive T-cells which have received cognate antigen signalling using Tocky-Angle (*34*) and multidimensional Tocky analysis (*33*).

The immunisation of S1a or S2b induced S-reactive T-cell populations including CXCR5^+^PD-1^hi^ Tfh cells. The reduction in the percentage of Timer+ cells in the Tfh fraction upon challenge after a single dose of S1a immunisation suggested that S1a-reactive Tfh cells vigorously proliferated to lose Timer protein. Recently, Merkenschlager et al showed that, using a fluorescent ubiquitin-based cell cycle indicator (Fucci) (*41*), Tfh cells enter cell cycle, showing vigorous proliferation in the first week after immunisation, as assessed by EdU and Ki67 staining, with 10-20% of Tfh cells EdU^+^ (*42*). Importantly, the repeated S1a immunisation induced not only Tfh but also associated transitional populations including Blue^+^ CD44^hi^ Tconv, indicating their proliferation and expansion as populations.

It is likely that the repeated doses of S1a immunisation expanded CD44^hi^ memory-phenotype Tconv, which promptly respond to the challenge dose of S1a, showing Blue fluorescence. This suggests that the microenvironment in the lymph node does not inhibit antigen presentation and subsequent TCR signalling in T-cells. Therefore, it is an intriguing finding that, after S1a boost immunisation, the majority of PD-1^hi^ CXCR5^+^ Tfh (∼80%) showed Timer Angle in the Arrested locus and the rest were mostly in the PAt locus (Fig. 3H). This indicates that the majority of the Tfh population no longer persistently engage with the antigen and only received intermittent TCR signals. This is compatible with the microscopic finding by Shulman et al that each of a stable B-Tfh interaction occurs between 5 – 30 min only (*43*). It is also possible that PD-1 on the S1a-reactive Tfh cells interacts with its ligands and inhibits TCR signalling through recruitment of the phosphatase SHP2 (*44*).

Shi et al reported that, since PD-L1 is constitutively expressed on follicular naïve B-cells, PD-1^hi^ CXCR5^+^ Tfh do not succumb to PD-1 mediated suppression inside the GC but do so in the mantle zone, which controls the positioning of Tfh cells inside the GC (*6*). While Tocky provides insights into T-cell dynamics within a unique time window, independently from microscopic studies, it does not provide spatial information. Future studies using the combination of Tocky and microscopy with appropriate lineage tracing markers are required to further address spaciotemporal dynamics of Tfh cells in vivo.

The findings on Tfh by us and others are relevant in COVID-19, as the infection as well as SARS-CoV-2 vaccines can induce antigen-reactive Tfh cells, contributing to the quality of neutralising antibodies (*45*). The percentage of circulating Tfh cells from Peripheral Blood Mononuclear cells (PMBCs) are correlated with neutralising antibodies in convalescent patients (*46*). Tfh cells promote B-cell maturation, which includes somatic hypermutation and class switching, and their differentiation is controlled by the transcription factor Bcl6 (*5*). T-cell-specific deletion of *Bcl6* using *Cd4-Cre Bcl6*^*fl*/fl^ mice leads to a severe reduction of CXCR5^+^PD-1^hi^ Tfh cells (*47, 48*). However, *Cd4-Cre Bcl6*^*fl*/fl^ mice can produce neutralising antibodies against S-protein and RBD-specific antibodies, while Blc6 is required for somatic hypermutation into S-specific antibodies (*48*). In addition, our study has shown that S1a (which contains NTD) and S2a can induce Tfh. On the other hand, S1b (which contains RBD) increased activated Treg and also induced some degree of activation in Tfh, albeit it did not expand Tfh population. These results are compatible with antigen mapping in mice (*27*). Therefore, our study provides unique insights into the effects of ‘non-immunogenic’ antigens on the activation of CD4 T-cells and Treg. Future studies using scanning peptides will further characterise the properties of each of T-cell epitopes in SARS-CoV-2 proteins.

Our analyses using the two Tocky models have shown that S1b-reactive CD25^hi^GITR^hi^PD-1^int^ Treg intermittently received TCR signalling and sustained *Foxp3* transcription over time. Persistent Foxp3 transcription is a hallmark of activated Treg with enhanced suppressive activities, or effector Treg (*35*). *Foxp3* transcription can be induced by TCR signalling, while it remains to be elucidated if TCR signalling drives effector Treg differentiation (*40*). CD25 and GITR are controlled by not only TCR signalling but also Foxp3 (*49*), and persistent Foxp3 transcription further upregulates CD25 and GITR expression in effector Treg (*35*). On the other hand, it is interesting that the S1b-reactive Treg have an intermediate PD-1 expression only. PD-1 (*Pdcd1*) transcription is activated by NF-κB and NFAT, which are downstream of TCR signalling (*50*). The intermediate expression of PD-1 suggests that S1b-reactive CD25^hi^GITR^hi^PD-1^int^ Treg succumb to PD-1-mediated suppression, which can decrease their PD-1 expression (*51*). Our combined approach using Foxp3 fate-mapping and Foxp3-Tocky showed that YFP+ cells had lower cells in the Persistent and PAt loci, which are enriched with CD25^hi^GITR^hi^PD-1^int^ Treg, in comparison to control. This indicates that S1b-reactive Treg, as identified by Foxp3-Tocky, are not found in the YFP+ fraction, which is composed of pre-existing thymic Treg that are reactive to self or microbiome antigens (*8, 12*), suggesting that S1b-reactive Treg are derived from non-Treg CD4 T-cells.

In addition, it remains to be addressed if S1b-reactive CD25^hi^GITR^hi^PD-1^int^ Treg have any association to activated Treg in COVID-19 infection (*52*). Future studies are needed to investigate in vivo dynamics of S-protein-reactive Treg and their significance in protective immunity and infection, using an infection model and single cell technologies such as single cell RNA-sequencing.

## MATERIALS AND METHODS

### Mice

*Nr4a3*-Tocky mice, or bacterial artificial chromosome (BAC) transgenic (Tg) *(Nr4a3*^*ΔExon3 FTfast*^), were generated by the Ono group and reported previously (*34*). *Nr4a3*-Tocky carries the BAC clone including the *Nr4a3* gene, which first coding exon was replaced with the Fluorescent Timer (FTfast(*53*)) gene using a knock-in knock-out approach. Nr4a3-Tocky mice were bred with *Foxp3*-IRES-GFP (C.Cg-Foxp3tm2Tch/J, Jax ID 006769) and maintained in a C57BL/6 background. Foxp3-Tocky mice were generated in the Ono lab and was reported previously (*35*). The Foxp3-Tocky::Foxp3 ^EGFP-Cre-ERT2^::R26R-EYFP triple transgenic line was bred by crossing Foxp3-Tocky with Foxp3 ^EGFP-Cre-ERT2^ (Jax ID 016961) and *R26R-EYFP* (Jax ID 006148) (*36*). All animal experiments were approved by the Animal Welfare and Ethical Review Body at Imperial College London and all animal work was performed in compliance with Home Office and Animal Scientific Procedures Act 1986 in the UK.

### Immunisation

Mice were subcutaneously immunised in the flank with 1.5 μg of S-protein fragment emulsified in Freund’s Incomplete adjuvant (IFA) (Sigma) containing 50 μg Poly(I:C) (Sigma). Control mice and immunisation regimens are described along the text.

### Expression and purification of recombinant apyrases

The S protein fragments were produced using a yeast system. First, synthetic genes s1a, s1b, s2a, and s2b (Invitrogen) were cloned into pPICZa-A Pichia vector and transformed into Pichia pastoris X-33. Expression was optimised for each protein then scaled up following the EasySelect Pichia Expression protocol (Invitrogen). His-tagged proteins were purified from yeast supernatants by Ni-NTA resin affinity chromatography and protein concentration was determined using Coomassie (Bradford) Protein Assay Kit (Thermo Fisher Scientific).

Purified proteins used for in vivo purposes were cleared from pyrogens using high-capacity endotoxin removal spin columns (Piece, Thermo Fisher Scientific). Protein samples were analysed by Western blotting. Samples were resolved by SDS-12% polyacrylamide gel electrophoresis under standard conditions and transferred to a polyvinylidene difluoride (PVDF) membrane (Amersham Hybond, GE Healthcare) using the Trans-Blot turbo transfer system (Biorad). Membrane was blocked (5 % w/v skimmed milk powder), probed with mouse anti c-myc primary antibody (9E10, Thermo Fisher Scientific) then incubated with goat anti mouse Ig-horseradish peroxidase secondary antibody (Thermo Fisher Scientific). Protein bands were visualized using enhanced chemiluminescence (ECL) Western Blotting Detection Reagents (Amersham Bioscience). Chemiluminescence was detected using a LAS-3000 Fuji Imager.

### Fluorescence Activated Cell Sorting (FACS) analysis

Draining inguinal lymph nodes (DLN) were harvested and single-cell suspensions were prepared by mechanical disruption in a buffer containing 10% PBS (D8537, Sigma), 2% Fetal Bovine Serum (FBS) (F9665, Sigma)). Staining was performed in U-bottom 96-well plate (650101, Greiner). Analysis was performed in BD LSRFortessa III and Cytek Aurora. For all experiments, cells were first incubated with fixable eFluor 780 fluorescent viability dye (65-0865-14, Invitrogen) and Fc-block (14-0161-82, eBioscience). Next, cells were incubated with the following biotinylated antibodies: CD11b (clone M1/70, eBioscience), CD11c (clone N418, eBioscience), TCR-γδ (eBioGL3, eBioscience), CD19 (clone eBio1D3, eBioscience), NK1.1 (clone PK136, Biolegend), Ly6G/C (clone R86-8C5, eBioscience), NK1.1 (clone PK136, Biolegend) and TER-119 (clone TER-119, eBioscience). Then, cells were incubated with Streptavidin APC-Cy7 (Biolegend) together with directly conjugated antibodies from the following list: CD4 BUV395 (clone GK1.5; BD), CD8 BV785 (clone 53–6.7; BioLegend), TCRβ BUV737 (clone H57-597; BD), PD-1 APC (clone J43; Biolegend), CXCR5 PE-Cy7 (clone SPRCL5; eBioscience) or BV650 (clone L138D7; BioLegend), CD25 PerCP-Cy5.5 (clone PC61.5; eBioscience) or PE-Cy7 (PC61.5; Tombo Bioscience), Alexa Fluor 700 (clone IM7; BioLegend), CD69 PE (clone H1.2F3; eBioscience), GITR BV605 (clone DTA-1; BD) or BV650 (clone DTA-1; BD), OX40 BV711 (clone OX-86; BioLegend). For all analysis, cells were first gated as lymphocytes, single cells, TCRβ^+^ APC-Cy7^-^. For in vitro detection of S-protein fragment specific Timer-Blue expression, DLN cells were counted using magnetic beads and 1 × 10^6^ total DLN cells were incubated 48 hours at 37°C in RPMI 1640 (R8758, Sigma) containing 2% FBS and 0.1% Penicillin-Streptomycin (P4333, Sigma) in the corresponding S-protein fragment (1 μg/mL).

### Angle transformation and Tocky UMAP analysis

The method for multidimensional Tocky UMAP analysis is described elsewhere (*33*). Briefly, the threshold of Timer fluorescence was set using wild-type T-cells. The algorithms for Timer Angle transformation and Tocky Locus analysis were reported previously (*54*). UMAP analysis used Timer Angle and Intensity and marker expression data using the CRAN package *umap* (*55*). Heatmaps were generated by the CRAN package *ggplot2* (*56*). Computational clustering was performed using k-means clustering and PCA output, in which k was determined by Bayesian information criterion (BIC) of *mclust* (*57*).

### Statistical analysis

Statistical analysis was performed using Prism 6 (GraphPad Software). Significance differences were calculated using two-tailed Mann-Whitney test, parametric two-tailed t-test and two-way ANOVA., *p<0.05, **p<0.01, ***p<0.001. Where indicated, the data from each independent experiment were normalised relative to controls (each individual value from both control and experimental samples is divided by the average control value) and pooled.

## ACKNOWLEDGEMENTS

We would like to thank Dr. Jessica Rowley and Dr. Larissa Zárate-García from the Flow Cytometry Facility South Kensington and the CBS team (mouse husbandry) of the Imperial College London. MO was supported by a CRUK Programme Foundation Award and the MRC grant (MR/S000208/1) and KAKENHI research grants from the Japan Society for the Promotion of Science (JSPS) (JP19H05426, JP21K07082, and JP21H00433).

## REFERENCES

1. P. Moss, The T cell immune response against SARS-CoV-2. Nature Immunology 23, 186–193 (2022).

2. A. Casrouge et al., Size estimate of the alpha beta TCR repertoire of naive mouse splenocytes. J Immunol 164, 5782–5787 (2000).

3. K. Kedzierska et al., Quantification of repertoire diversity of influenza-specific epitopes with predominant public or private TCR usage. J Immunol 177, 6705–6712 (2006).

4. L. L. Pewe, J. M. Netland, S. B. Heard, S. Perlman, Very diverse CD8 T cell clonotypic responses after virus infections. J Immunol 172, 3151–3156 (2004).

5. S. Crotty, Follicular helper CD4 T cells (TFH). Annu Rev Immunol 29, 621–663 (2011).

6. J. Shi et al., PD-1 Controls Follicular T Helper Cell Positioning and Function. Immunity 49, 264–274 e264 (2018).

7. J. C. Sun, M. J. Bevan, Defective CD8 T cell memory following acute infection without CD4 T cell help. Science 300, 339–342 (2003).

8. E. V. Russler-Germain, S. Rengarajan, C. S. Hsieh, Antigen-specific regulatory T-cell responses to intestinal microbiota. Mucosal Immunology 10, 1375–1386 (2017).

9. A. P. Basto, L. Graca, Regulation of antibody responses against self and foreign antigens by Tfr cells: implications for vaccine development. Oxford Open Immunology 2, (2021).

10. P. T. Sage, A. H. Sharpe, T follicular regulatory cells. Immunological Reviews 271, 246–259 (2016).

11. S. Kumar et al., Developmental bifurcation of human T follicular regulatory cells. Sci Immunol 6, (2021).

12. M. S. Jordan et al., Thymic selection of CD4+CD25+ regulatory T cells induced by an agonist self-peptide. Nat Immunol 2, 301–306 (2001).

13. J. T. Jacobsen et al., Expression of Foxp3 by T follicular helper cells in end-stage germinal centers. Science 373, (2021).

14. J. J. Moon et al., Tracking epitope-specific T cells. Nat Protoc 4, 565–581 (2009).

15. M. L. Bettini, M. Bettini, D. A. Vignali, T-cell receptor retrogenic mice: a rapid, flexible alternative to T-cell receptor transgenic mice. Immunology 136, 265–272 (2012).

16. K. A. Hogquist, S. C. Jameson, The self-obsession of T cells: how TCR signaling thresholds affect fate ‘decisions’ and effector function. Nature Immunology 15, 815–823 (2014).

17. A. Sacchetti, T. El Sewedy, A. F. Nasr, S. Alberti, Efficient GFP mutations profoundly affect mRNA transcription and translation rates. FEBS Lett 492, 151–155 (2001).

18. R. J. Martinez, R. Andargachew, H. A. Martinez, B. D. Evavold, Low-affinity CD4+ T cells are major responders in the primary immune response. Nature Communications 7, 13848 (2016).

19. T. Maruhashi et al., LAG-3 inhibits the activation of CD4(+) T cells that recognize stable pMHCII through its conformation-dependent recognition of pMHCII. Nat Immunol 19, 1415–1426 (2018).

20. Y. Peng et al., Broad and strong memory CD4+ and CD8+ T cells induced by SARS-CoV-2 in UK convalescent individuals following COVID-19. Nature Immunology 21, 1336–1345 (2020).

21. U. Sahin et al., BNT162b2 vaccine induces neutralizing antibodies and poly-specific T cells in humans. Nature 595, 572–577 (2021).

22. D. Martínez-Flores et al., SARS-CoV-2 Vaccines Based on the Spike Glycoprotein and Implications of New Viral Variants. Frontiers in Immunology 12, (2021).

23. W. T. Harvey et al., SARS-CoV-2 variants, spike mutations and immune escape. Nature Reviews Microbiology 19, 409–424 (2021).

24. F. Schmidt et al., High genetic barrier to SARS-CoV-2 polyclonal neutralizing antibody escape. Nature 600, 512–516 (2021).

25. A. Grifoni et al., SARS-CoV-2 human T cell epitopes: Adaptive immune response against COVID-19. Cell Host Microbe 29, 1076–1092 (2021).

26. A. Tarke et al., Impact of SARS-CoV-2 variants on the total CD4^+^ and CD8^+^ T&#xa0;cell reactivity in infected or vaccinated individuals. Cell Reports Medicine 2, (2021).

27. Z. Zhuang et al., Mapping and role of T cell response in SARS-CoV-2–infected mice. Journal of Experimental Medicine 218, (2021).

28. S. R. Leist et al., A Mouse-Adapted SARS-CoV-2 Induces Acute Lung Injury and Mortality in Standard Laboratory Mice. Cell 183, 1070–1085 e1012 (2020).

29. N. Iwata-Yoshikawa et al., A lethal mouse model for evaluating vaccine-associated enhanced respiratory disease during SARS-CoV-2 infection. Sci Adv 8, eabh3827 (2022).

30. B. Kingstad-Bakke et al., Vaccine-induced systemic and mucosal T cell immunity to SARS-CoV-2 viral variants. Proc Natl Acad Sci U S A 119, e2118312119 (2022).

31. S. Bortoluzzi et al., Brief homogeneous TCR signals instruct common iNKT progenitors whose effector diversification is characterized by subsequent cytokine signaling. Immunity 54, 2497-2513. e2499 (2021).

32. G. Bozhanova et al., CD4 T cell dynamics shape the immune response to combination oncolytic herpes virus and BRAF inhibitor therapy for melanoma. Journal for immunotherapy of cancer 10, (2022).

33. J. Hassan et al., Single-cell level temporal profiling of tumour-reactive T cells under immune checkpoint blockade. bioRxiv, 2022.2007.2019.500582 (2022).

34. D. Bending et al., A timer for analyzing temporally dynamic changes in transcription during differentiation in vivo. J Cell Biol 217, 2931–2950 (2018).

35. D. Bending et al., A temporally dynamic Foxp3 autoregulatory transcriptional circuit controls the effector Treg programme. The EMBO journal 37, e99013 (2018).

36. Y. P. Rubtsov et al., Stability of the regulatory T cell lineage in vivo. Science 329, 1667–1671 (2010).

37. S. Srinivas et al., Cre reporter strains produced by targeted insertion of EYFP and ECFP into the ROSA26 locus. BMC Dev Biol 1, 4 (2001).

38. B. J. Meckiff et al., Imbalance of Regulatory and Cytotoxic SARS-CoV-2-Reactive CD4(+) T Cells in COVID-19. Cell 183, 1340–1353 e1316 (2020).

39. S. Z. Josefowicz, L. F. Lu, A. Y. Rudensky, Regulatory T cells: mechanisms of differentiation and function. Annu Rev Immunol 30, 531–564 (2012).

40. M. Ono, Control of regulatory T-cell differentiation and function by T-cell receptor signalling and Foxp3 transcription factor complexes. Immunology 160, 24–37 (2020).

41. A. Sakaue-Sawano et al., Visualizing Spatiotemporal Dynamics of Multicellular Cell-Cycle Progression. Cell 132, 487–498 (2008).

42. J. Merkenschlager et al., Dynamic regulation of TFH selection during the germinal centre reaction. Nature 591, 458–463 (2021).

43. Z. Shulman et al., Dynamic signaling by T follicular helper cells during germinal center B cell selection. Science 345, 1058–1062 (2014).

44. T. Okazaki, A. Maeda, H. Nishimura, T. Kurosaki, T. Honjo, PD-1 immunoreceptor inhibits B cell receptor-mediated signaling by recruiting src homology 2-domain-containing tyrosine phosphatase 2 to phosphotyrosine. Proc Natl Acad Sci U S A 98, 13866–13871 (2001).

45. K. Lederer et al., SARS-CoV-2 mRNA Vaccines Foster Potent Antigen-Specific Germinal Center Responses Associated with Neutralizing Antibody Generation. Immunity 53, 1281-1295.e1285 (2020).

46. S. Boppana et al., SARS-CoV-2-specific circulating T follicular helper cells correlate with neutralizing antibodies and increase during early convalescence. PLOS Pathogens 17, e1009761 (2021).

47. K. Hollister et al., Insights into the role of Bcl6 in follicular Th cells using a new conditional mutant mouse model. J Immunol 191, 3705–3711 (2013).

48. J. S. Chen et al., High-affinity, neutralizing antibodies to SARS-CoV-2 can be made without T follicular helper cells. Science Immunology 7, eabl5652.

49. S. Hori, T. Nomura, S. Sakaguchi, Control of Regulatory T Cell Development by the Transcription Factor Foxp3. Science 299, 1057–1061 (2003).

50. A. P. Bally, J. W. Austin, J. M. Boss, Genetic and Epigenetic Regulation of PD-1 Expression. J Immunol 196, 2431–2437 (2016).

51. H. Okamura et al., PD-1 aborts the activation trajectory of autoreactive CD8(+) T cells to prohibit their acquisition of effector functions. J Autoimmun 105, 102296 (2019).

52. B. Kalfaoglu, J. Almeida-Santos, C. A. Tye, Y. Satou, M. Ono, T-Cell Hyperactivation and Paralysis in Severe COVID-19 Infection Revealed by Single-Cell Analysis. Front Immunol 11, 589380 (2020).

53. F. V. Subach et al., Monomeric fluorescent timers that change color from blue to red report on cellular trafficking. Nat Chem Biol 5, 118–126 (2009).

54. D. Bending et al., A timer for analyzing temporally dynamic changes in transcription during differentiation in vivo. Journal of Cell Biology 217, 2931–2950 (2018).

55. T. Konopka. (https://CRAN.R-project.org/package=umap 2022).

56. H. Wickham, ggplot2: Elegant Graphics for Data Analysis. (Springer-Verlag New York, 2016).

57. L. Scrucca, M. Fop, T. B. Murphy, A. E. Raftery, mclust 5: Clustering, Classification and Density Estimation Using Gaussian Finite Mixture Models. R J 8, 289–317 (2016).

