## Supplementary Figure Legends for "Temporal profiling of CD4 T-cell activation and differentiation upon SARS-CoV-2 spike protein immunisation"

**Figure S1. The effects of S1a immunisation on antigen-reactive CD4 T-cells**

Nr4a3-Tocky mice were immunised with either S1a fragment or PBS emulsified in Incomplete Freund’s adjuvant (IFA) containing 50µg of Poly (IC) and challenged 7 days later in absence of added adjuvant. Analysis took place 24h later. (**A-C)** Shown are the expression of Nr4a3-Blue and -Red and the percentage of Nr4a3 Timer^+^ cells in the total CD4 population (A), the expression of Foxp3 and percentage of Foxp3^+^ cells within the CD4 population (B), and the percentage of Nr4a3 Timer^+^ cells in CD4 cells decomposed into Tconv and Treg (C). (**D**) Shown are the normalised Timer intensity data in New, NPt and Persistent Timer locus of Tconv PD1^hi^ CXCR5^+^ cells. Data from two independent experiments (N=4-5). (**E)** Uniform Manifold Approximation and Projection (UMAP) analysis of Tconv Timer^+^ cells from immunised Nr4a3-Tocky mice. Heatmaps show Timer Angle and Intensity as well as the expression of the indicated markers. The UMAP was applied to Tconv Timer^+^ cell data from draining lymph nodes (DLN) of both S1a and PBS immunised mice. UMAP was performed using Timer-Blue, Timer-Red, CD25, CD44, GITR, PD1, CXCR5, and CD69 as input data. **(F)** Distribution of each individual Cluster identified by k-means clustering in the UMAP space from (E). (**G)** Percentage of Tconv Timer^+^ cells in each Cluster identified in (F) from either S1a or PBS samples (n=4). **(H-J)** UMAP and k-means clustering was performed in Timer^+^ Treg cells. Shown are heatmaps of Timer Angle and Intensity as well as the expression of the indicated markers (H), the distribution of each individual Cluster identified by k-means clustering in the UMAP space (I), and the percentage of Treg Timer^+^ cells in each Cluster from samples of both groups (J) (N=4). Statistical analysis performed using nonparametric and parametric T-test *P < 0.05, **P < 0.005.

**Figure S2. The effects of a booster S1a immunisation on antigen-reactive Tfh cells and transitional population.** Nr4a3-Tocky mice were immunised with 1.5µg of S1a fragment and after two weeks received either boost S1a (Boost) or PBS (Single). Two weeks later both groups were challenged with S1a and analysis took place 24h later. **(A)** Percentage of total Nr4a3-Timer^+^ cells in both total CD4 (Left), Tconv (Middle) and Treg cells (Right). **(B)** UMAP analysis was performed using Tconv Timer^+^ cell data and shown are the individual Clusters identified by k-means clustering overlaid in the UMAP space. **(C)** Shown are the percentages of Tconv Timer^+^ cells in each cluster identified in (B) except Clusters 4, 6 and 9. **(D)** Violin plots showing Timer Intensity as well as the expression of the indicated markers in each Timer locus from Clusters 4, 6 and 9 as well as all concatenated Clusters from Figure 3E. Data from one experiment (N=3).

**Figure S3. The effects of S1b immunisation on antigen-reactive CD4 T cells and Treg.** **(A)** Nr4a3-Tocky mice were immunised with either S1b fragment or PBS emulsified in Incomplete Freund’s adjuvant (IFA) containing 50µg of Poly (IC) and challenged 7 days later in absence of added adjuvant. Analysis took place 24h later and shown is the percentage of Nr4a3-Timer^+^ Tconv cells. Data from 2 independent experiments (N=2-4). **(B-D)** Uniform Manifold Approximation and Projection (UMAP) analysis of Tconv Timer^+^ cells from mice in (A). The UMAP was performed using Timer-Blue, Timer-Red, CD25, CD44, GITR, PD1, CXCR5, and CD69 as input data. Shown are heatmaps of Timer Angle and Intensity as well as expression of the indicated markers (B), the distribution of Clusters identified by k-means clustering in the UMAP space (C), and the percentage of Tconv Timer^+^ cells in the indicated Clusters in samples from both groups (N=4) (D). **(E-F)** Analysis of CD4 Foxp3^+^ PD1^hi^ CXCR5^+^ cells from mice immunised in (A). Shown are their normalised percentage of total Timer^+^ cells (E) and their expression of Nr4a3-Blue and –Red (F). Data from 2 independent experiments (N=2-4). **(G-H)** UMAP and k-means clustering analysis of Treg Timer^+^ cells from mice immunised as in (A). Shown are the identified Clusters overlaid in the UMAP space (G) and the percentage of Treg Timer^+^ cells in each of the identified Clusters (H). **(I)** Mice were immunised with S1b and after two weeks received either boost S1b immunisation or PBS in absence of added adjuvant. Two weeks later all mice were challenged with S1b in absence of added adjuvant and draining lymph nodes were analysed 24h later. **(J)** Shown are the normalised percentage of Foxp3^-^ PD1^hi^ CXCR5^+^ (Left) and Foxp3^+^ PD1^hi^ CXCR5^+^ (Right) in total CD4 T cells from mice immunised as in (G) (N=3). Statistical analysis performed using nonparametric T-test *P < 0.05, **P < 0.005.

**Figure S4. The effects of S2a and S2b immunisation on antigen-reactive CD4-T cells and Treg. (A)** Nr4a3-Tocky mice were immunised with either S2a fragment or PBS emulsified in Incomplete Freund’s adjuvant (IFA) containing 50µg of Poly (IC) and challenged 7 days later in absence of added adjuvant. Analysis took place 24h later. **(B-D)** Shown are the normalised percentage of Tconv PD1^hi^ CXCR5^+^ in CD4 T cells (B), their expression of Nr4a3-Blue and –Red (C) and the normalised percentage of Nr4a3-Timer^+^ cells in Tconv PD1^hi^ CXCR5^+^ (Left) total Tconv (Middle) and Treg (Right) cells from mice in (A) (D). Data from 3 independent experiments (N=2-4). **(E-H)** UMAP and k-means clustering of Tconv and Treg cells from mice immunised as in (A). Shown are the distribution of each identified Cluster in the Tconv UMAP space and the percentage of cells in each cluster (E-F), and the distribution of each identified Cluster in the Treg UMAP space and the percentage of cells in each cluster (G-H) (N=4). **(I-J)** Mice were immunised with either S2b fragment or PBS emulsified in Incomplete Freund’s adjuvant (IFA) containing 50µg of Poly (IC) and challenged 7 days later in absence of added adjuvant. Analysis took place 24h later. Shown are the normalised percentage of Nr4a3-Timer^+^ cells in Tconv (I) and Treg (J) cells. Data from two independent experiments (N=2-4). **(K-L)** UMAP and k-means clustering analysis of Tconv Timer+ cells from mice immunised as in (I-J). Shown are the identified Clusters in the Tconv Timer^+^ UMAP space (K) and the percentage of Tconv Timer^+^ cells in each identified Cluster (L) except Cluster 1 (N=4). (**M-N**) UMAP and k-means clustering analysis of Treg Timer^+^ cells from mice immunised as in (I-J). Shown are the identified Clusters in the Treg Timer^+^ UMAP space (M) and the percentage of Treg Timer^+^ cells in each identified Cluster (N) (N=4). Statistical analysis performed using nonparametric T-test *P < 0.05, **P < 0.005.
