## Supplementary figures and images for "Temporal profiling of CD4 T-cell activation and differentiation upon SARS-CoV-2 spike protein immunisation"

### Supplementary Figure S1

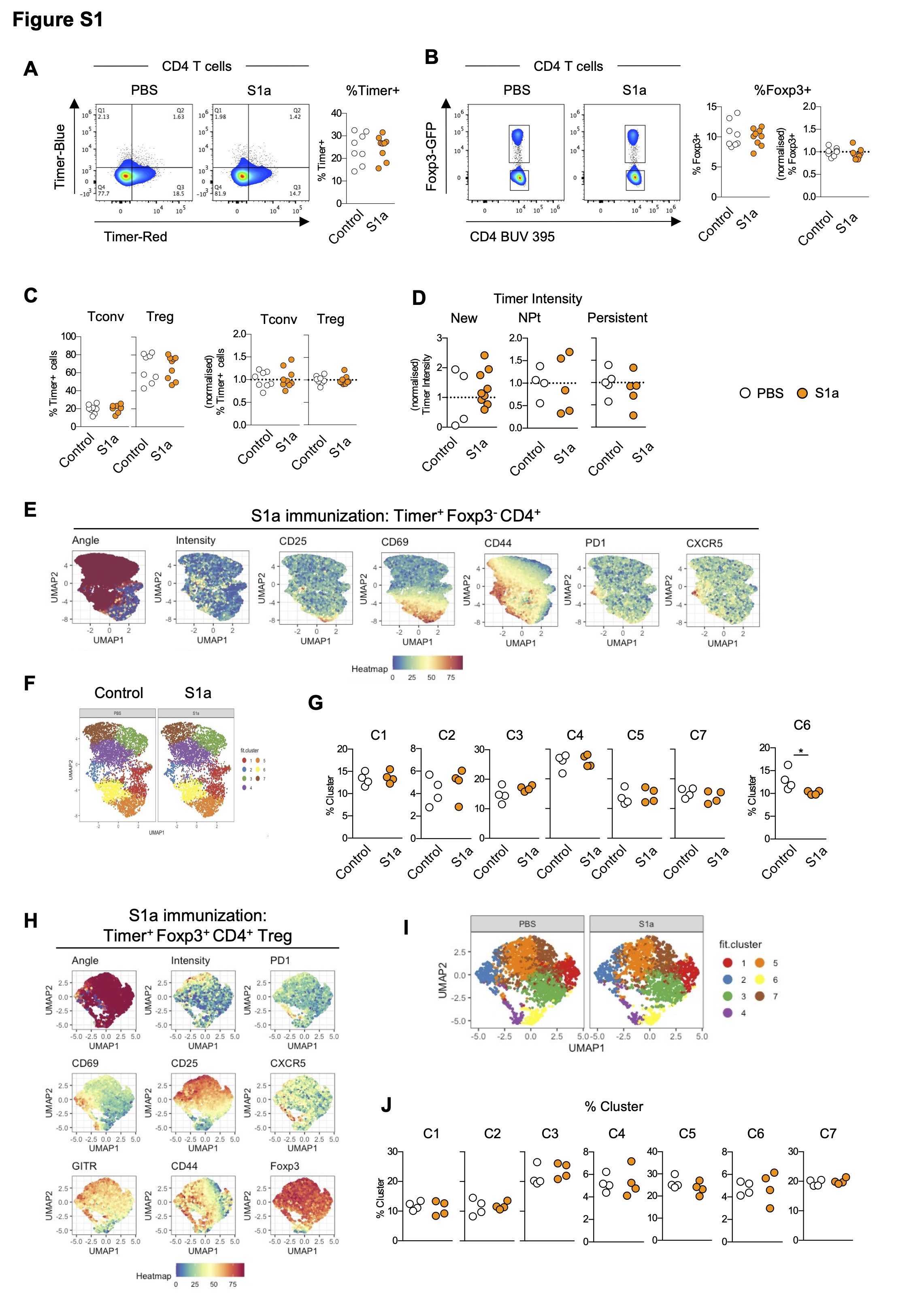

### Supplementary Figure S2

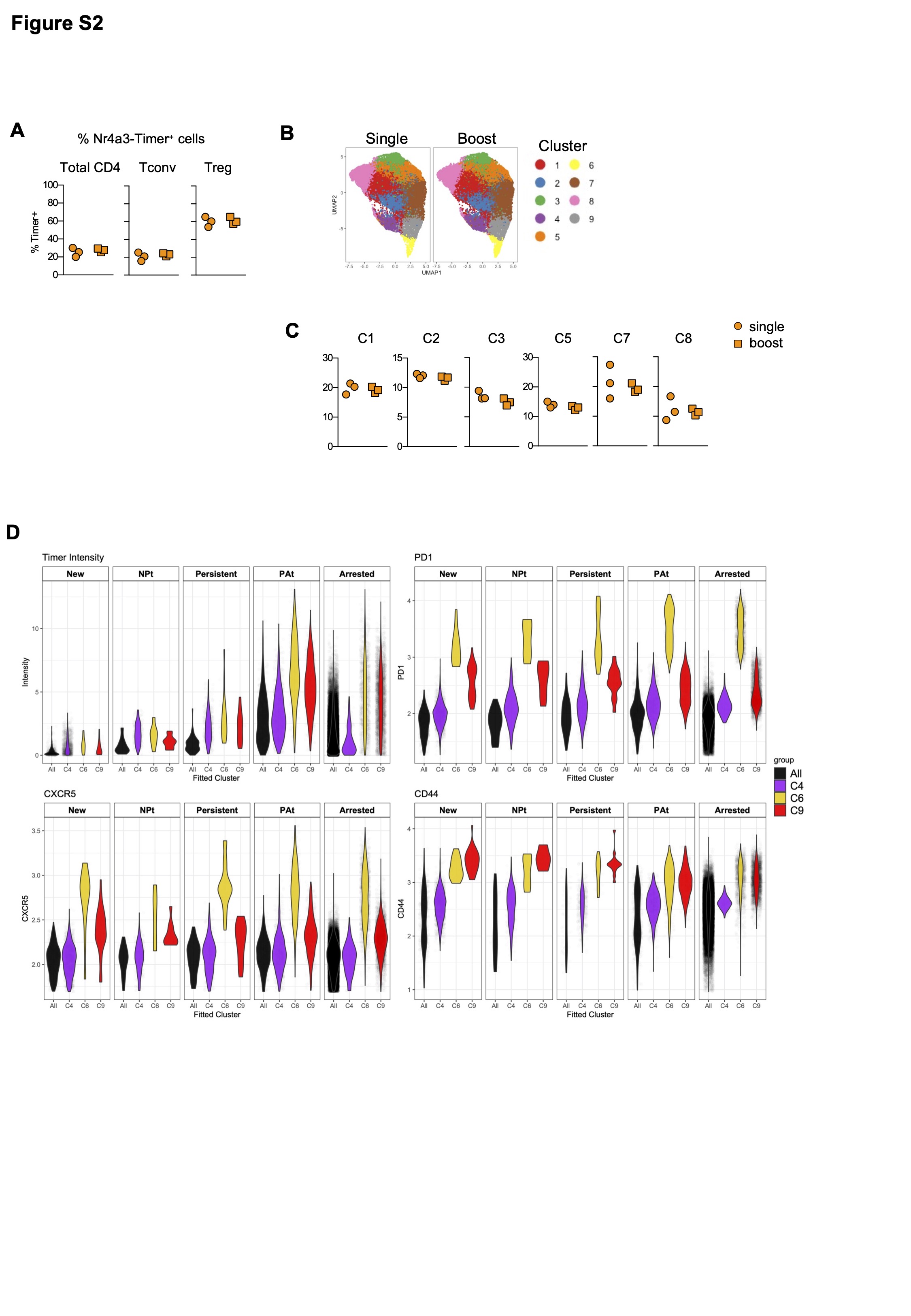

### Supplementary Figure S3

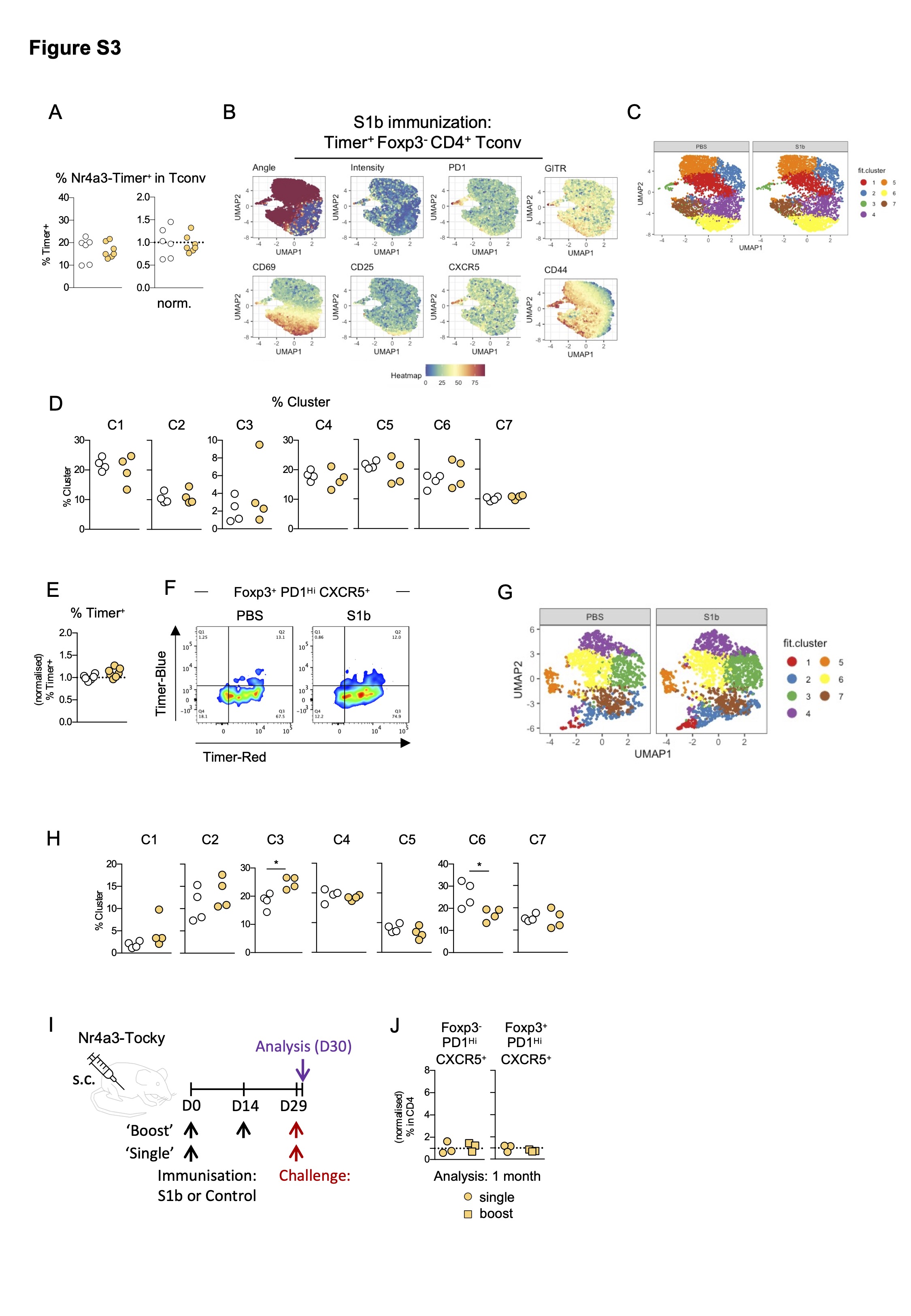

### Supplementary Figure S4

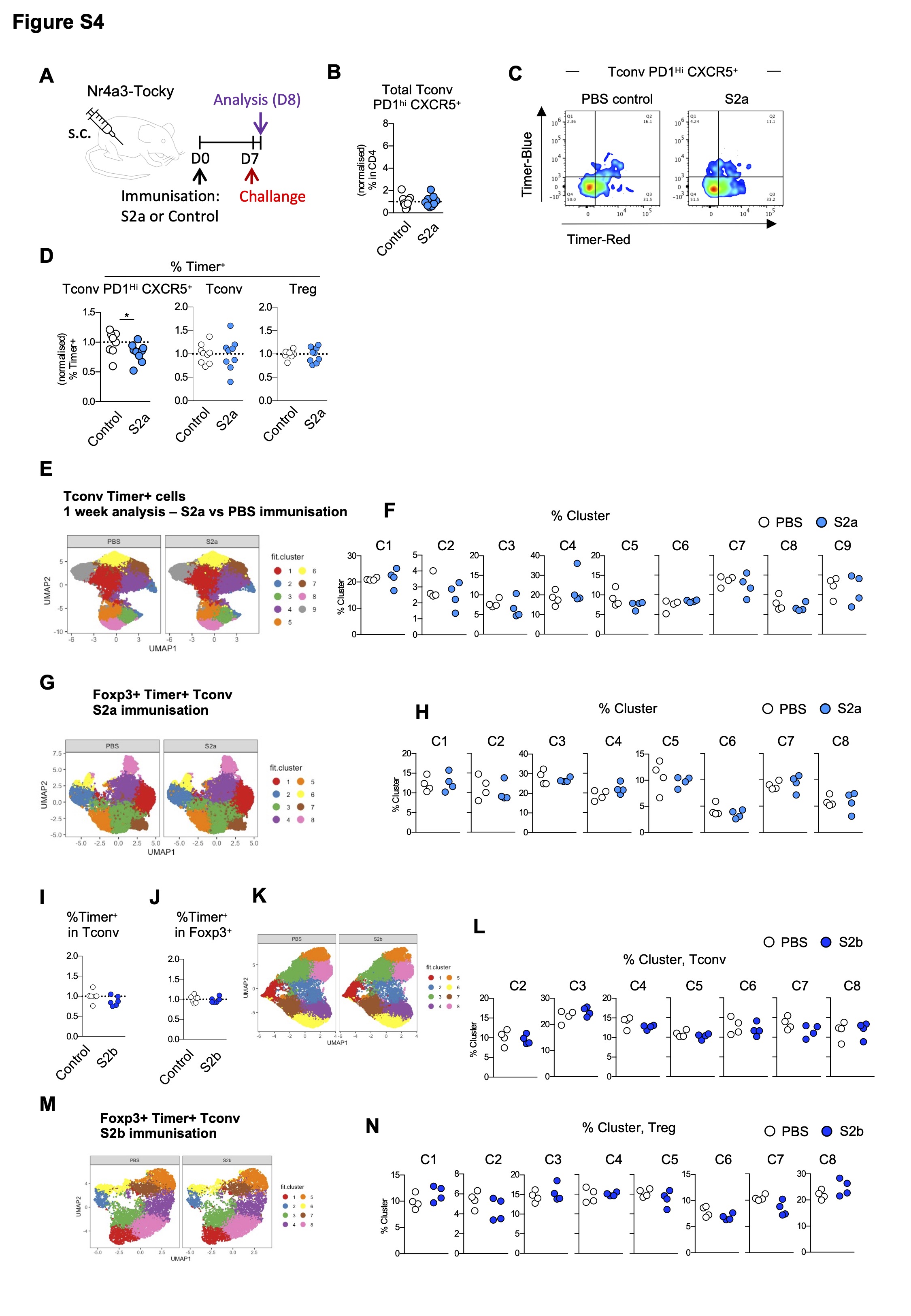
